# Science Podcasts: Analysis of Global Production and Output From 2004 to 2018

**DOI:** 10.1101/298356

**Authors:** Lewis E. MacKenzie

## Abstract

Since 2004, podcasts have emerged as a decentralised medium for science communication to the global public. However, to-date, there have been no large-scale quantitative studies of the production and dissemination of science podcasts. This study identified 952 English language science podcasts available between January and February 2018 and analysed online textual and visual data related to the podcasts and classified and noted key production parameters. It was found that the total number of science podcast series available grew linearly between 2004 and 2010, and then exponentially between 2010 and 2018. 65% of science podcast series were hosted by scientists and 77% were targeted to public audiences. Although a wide range of primarily single-subject science podcasts series were noted, 34% of science podcast series were not dedicated to a science subject. Compared to biology and physics, chemistry may be under-represented by science podcasts. Only 24% of science podcast series had any overt financial income. 62% of science podcast series were affiliated to an organisation; producing a greater number of episodes (median = 24, average = 96) than independent science podcast series (median = 16, average = 48). This study provides the first ‘snapshot’ of how science podcasts are being used to communicate science to public audiences around the globe.

## 2. Introduction

Since 2004, podcasts have emerged as a new decentralised medium for free and independent communication to global audiences. Podcasts are typically audio-only, hosted online, and distributed to audiences via direct, on-demand audio and video downloads to personal computers, MP3 players, interactive media devices, and smartphones.[1] For app-enabled devices, episodes of a podcast series can be automatically downloaded via free opt-in subscription to particular podcast series ‘feeds’.[2]^a^ For audiences, audio podcasts are particularly convenient because they can be listened-to whilst undertaking other activities without looking at a screen. Additionally, podcast may be accompanied supplementary ‘show notes’ that can contain text, hyperlinks, and/or images. For content creators, podcasts are convenient because they can be created with readily-available equipment, i.e. a microphone, audio recording/editing software, a web hosting service,[6] or even just a single smartphone.[7] Despite minimal technical requirements, podcasts can also be created with high-end professional production values, similar to broadcast radio shows.

Science podcasts have become a varied and abundant avenue for science communication, with many hundreds of English language science podcast series currently available to the public, covering many different topics, audiences, and formats. Due to being unconstrained by the format demands of TV and radio media, many diverse styles of science podcasts are available, including: monologues, informal chats, professional science news, panel shows, and comedy.[8] The freedom to incorporate humorous elements (if desired) is particularly notable because humour has been beneficial for engaging audiences in science communication.[9,10] Crucially, podcasts enable science communicators to directly engage audiences in a style of their choosing, without the risks of miscommunication associated with “stage managed” dissemination via traditional print and broadcast media.[11]

Due to their online distribution, podcasts have the potential to reach audiences around the globe, in a manner unconstrained by the demographic or geographic restrictions associated with traditional regional or national media.[12] This allows some podcasts to cater for niche audiences that are not a priority for traditional media. One such example of a highly specialised science podcast series is: ‘*This Week in Virology*’, which primarily serves the virology research community, yet which also reportedly has a large proportion of public listeners.[9] Another example of podcasts filling an under-served niche are podcasts that focus on science for young children, one example of which is *‘Wow In The World’*.[13]. Due to the large number of science podcasts, their accessible nature, and their varied production, it could be said that *“there is a science podcast for everyone”*.

For science communication. the audio-only format of podcasts provides several key advantages over traditional print and televisual media beyond that of convenience to listener and producer. Merzagora notes that compared to television and print, audio media is *“more relaxed and reflective”;* that it *“allows the audience to hear the true voice of the protagonist”* (i.e. the science communicator); and that *“the barrier separating the listener from journalists and scientists is less impenetrable”*.[14] Additionally, podcasts creators commonly use websites and social media to receive listener feedback and facilitate discussion. Such “two-way dialogue” – not typically available in traditional broadcast and print media - can help improve public trust in science”.[15,16] It has been speculated that podcast audiences may feel more personally connected to the producers of podcasts than of other forms of media.[17] Additionally, podcasts have been demonstrated to improve scientific information uptake in students, medical patients, and the public.[18–20] These advantages combine to make podcasts an attractive medium for science communication for both independent science communicators and larger organisations. Examples of large organisations with science podcasts include: professional scientific societies, space agencies, funding agencies/charities, scientific journals, government agencies, schools, and universities.

Audience engagement metrics for podcasting are either not well developed or not publicly available.[21] Therefore, studies of podcast listener demographics have primarily relied on audience surveys. In 2018, a commercial survey of general podcast audiences in the USA found that both men and women listen to podcasts in similar proportions (27% and 24% of respondents respectively); that podcast audiences skew towards young adults; that podcast audiences are well-educated, and that individuals typically listen to an average of 7 podcasts per week (corresponding to an average of 6 hours 37 minutes).[22] In contrast, a study of science podcast audiences in Brazil by Dantas-Queiroz et al.[10] found that an overwhelming proportion (87%) of self-reported responders to a science podcast survey were men; this may reflect wider societal biases influencing differences in how men and women engage with scientific content online,[10] but the constituent demographics of science podcast audiences are still unclear.

Despite the rise of podcasts as a popular medium for science communication, there have been no studies of the large-scale patterns in the production of science podcasts; this represents a large and fundamental gap in our knowledge of science communication. Therefore, this aim of this study was to provide the first large-scale quantitative insight into the overall global production and dissemination of science podcasts. This has been achieved by analysing online textual and visual presence of 952 English language science podcasts for key production variables, including: audio/visual format, topic, target audiences, hosts, number of episodes released, lifespan of podcasts, supplementary income, and the incorporation of supplementary show notes. All data associated with this study is available as a supplementary dataset in the form of a Microsoft Excel spreadsheet.

## 3. Materials and Methods

### Information Sources

All information used in this study was sourced from public websites that were dedicated to the promotion of podcasts. Information was gleaned exclusively from visual and textual “metadata” relating to each podcast series, including the description of each podcast series on ‘*iTunes’*, the websites of podcasts, and the social media content associated with podcast series, i.e. on *‘Twitter’*,[23]’*Facebook’,[24]* and *‘Patreon’.[25]*. The audio and video content of podcasts themselves was not utilized due to the impracticalities associated with listening and transcribing the tens of thousands of hours of audio content that science podcasts provide.[26] Producers and other individuals associated with the production of podcast series were not contacted for information relating to this study in order to avoid methodical disparity between podcast series with responsive producers and those without responsive producers. In all cases, information was accessed between the 5^th^ of January 2018 and 5^th^ of February 2018. The associated supplementary database contains all the specific dates of when each website URL was accessed. All data was manually coded and categorised the author.

### Identification of Podcast Series

Due to the decentralised nature of the podcast medium, there is not a single podcast database or website that lists all podcast series. However, the closest thing to a “de-facto” centralised podcast series database is the *‘iTunes’* podcast directory, which as of 2015, was estimated to list over 200,000 podcast series.[27]^b^ The *‘iTunes’* podcast directory’s search function is available cross-platform: i.e. it can be used by podcast apps running on non-Apple platforms, e.g. Android devices.[28,29] If a podcast series is not listed on the *‘iTunes’* podcast directory, then it is considerably less likely to be found by listeners.[30] Therefore, in line with other studies,[15] the *‘iTunes’* podcast directory was selected as the primary directory from which to source podcasts.

A systematic review of the *‘iTunes’* podcasts *‘Natural Sciences’* directory was conducted to identify potential podcast series for inclusion in this study.[31] All podcast series in the *‘Natural Sciences’* section were examined between the 5^th^ of January 2018 and the 5^th^ of February 2018 by proceeding through the section in reverse alphabetical order. However, it should be noted that the category a podcast series is assigned to within the *‘iTunes’* podcast directory is based entirely on the category nominated by the uploader of said podcast series [30]: consequently, there are many non-scientific podcast series spuriously listed in the *‘Natural Sciences’* ‘*iTunes*’ category.[31] Therefore, to ensure that only valid podcast series covering scientific topics were examined in this study, a stringent set of inclusion criteria were developed and applied (see sub-section ‘Categorical Definitions’). The inclusion criteria were applied after analysis of the textual and visual information associated with each podcast series and are defined in the sub-section ‘Inclusion/Exclusion Criteria’. Additionally, during the study, some podcast series were found that were not listed on the ‘*iTunes*’ podcast directory. These were also considered for inclusion. Of these *‘non-iTunes’* listed podcasts, 18 met the inclusion criteria, representing ~2% of the 952 science podcast series included in this study.

### Inclusion/Exclusion Criteria

To ensure that only legitimate science podcast series were included in this study, the following set of inclusion/exclusion criteria were developed and applied:

- Only English language podcast series were included in this study. If a podcast series was available in multiple languages, then only the English language podcast feed was analysed to avoid duplicating content.
- For the purposes of this study, “science podcasts” are primarily defined as podcast series covering topics in the natural sciences, i.e. physics, chemistry, biosciences, geology, oceanography, climate change, palaeontology, and mathematics. *Nb: this definition is functionally similar to that used by Birch and Weitkamp (2010).[15]*
- Under a secondary definition: podcast series covering the academic and research aspects of computer science, engineering, pharmacology and medicine were included. These podcast series account for 3% of the podcasts included in the study.
- Podcast series focusing on non-science topics were excluded. *Nb: examples of such topics include: consumer technology; business; gardening; bird-watching (“birding”); food/cooking; religion; life-coaching; weather; sustainability; environmental activism; pseudo-science; occult and paranormal; nerd culture, and podcasts primarily intended to review or sell commercial products, e.g. relating to tropical fish keeping or telescopes*.
- If the scientific nature of a podcast series was unclear, then that podcast series was excluded.
- If a podcast series was available as separate audio-only and video feeds covering the otherwise identical content, then only the video-feed was included for analysis to avoid data duplication.
- Podcast series with no episodes available to stream or download via either ‘*iTunes*’ or another website were excluded.
- To be included for analysis, episodes of a podcast series had to be freely available for listeners to stream or download from a source at the time of sampling. *For example, if a podcast had 100 episodes available on* ‘*iTunes’, yet had 250 episodes available to stream on their own website, then 250 episodes were noted*.
- If the content of a podcast series was originally available prior to 2004, (e.g. as an internet or broadcast radio show), then the original broadcast date of the first show episode was used in-lieu of the upload date of the podcast episode. *Nb: this was used because it provides some context for long-running internet radio series that have embraced the podcast format. However, this has some consequences for interpreting the results of this study: see the “Methodology and associated limitations” sub-section for more details*.

### Categorical Definitions

Podcast series, their production methods, and their production outputs were manually classified by the author in accordance with the definitions provided in Table 2 and the methods detailed herein.

Science podcast series were typically found to be focused on either a single distinct topic or to cover many different topics across a wide range of scientific disciplines. Therefore, an exclusive single-category system was used to classify the topics of podcast series; i.e. podcast series were either classified as a single subject, or if they covered many topics, they were classified as ‘general science’. Similarly, an exclusive one-category classification system was deemed sufficient for organisational affiliations, target audiences, and whether or not a podcast series was video or audio format. Three non-exclusive categories were devised for classifying supplementary income: *‘donations’, ‘merchandise’*, and *‘advertising/sponsorship’*. These categories were not exclusive as individual podcast series may employ some or all of these income mechanisms.

*‘Country of podcast production’* was defined as the country primarily associated with a podcast series and its hosts. For this category an exclusive, exclusive one-category classification system was adopted; if two or more countries were associated with a podcast series, then it was classed as ‘multinational’.

Science podcast hosts were classified according to a ranked classification system consisting of: *‘Scientific Researchers/Educators’* (Rank 5); *‘Media/Journalism Professionals’* (Rank 4); (3) *‘Other Professionals’* (Rank 3); *‘Amateurs’* (Rank 2); and *‘unclear’* (Rank 1), where the ranking is related to general expertise/ scientific authority, i.e. the higher the rank the higher the authority (see Table 2). In the case where podcasts had multiple hosts (or a single host of different areas of expertise) then the highest ranked category corresponding to one of the hosts was recorded, even if that host was in an overall minority of hosts. The limitations of this method are discussed in the ‘Methodology and associated limitations” sub-section of the discussion.

Podcast activity and podcast lifespans were determined by the objective definitions described in Table 2.

### Data analysis

All relevant information and resultant categorical analysis was recorded within a spreadsheet database (Microsoft Excel 2016, .xlsx format), which is available as a supplementary dataset to this manuscript. Basic categorical analysis was undertaken with Microsoft Excel, however, advanced categorical and data analysis (such as analysis of podcast series lifespan) was carried out using custom-written MATLAB scripts (MATLAB 2017b/ 2018a, Mathworks). Figures were created from data by plotting in MATLAB with some minor annotations added in PowerPoint (Microsoft PowerPoint 2016).

To estimate mean lifespan of podcast series, single-term and two-term exponential decays were fitted to podcast series lifespan data by least-squares regression.^c^ The equations describing these fits are respectively:

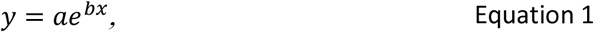

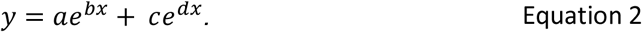

Where *a, b, c, and d*, are the recovered best-fit parameters with associated 95% confidence intervals. The mean lifespan (T) was then calculated by:

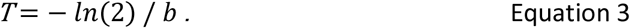

Where ln(2) is the natural logarithm of *2* (approximately 0.693). For estimation of long and short mean lifespans components from two-term exponential decay fits, *d* was substituted for *b* in Equation 3. 95% confidence intervals for the upper and lower bounds of T were also estimated. The statistical significance of the difference between the best-fit estimates of T for long duration and short duration components were estimated by the method described in Bland and Altman (2011), which is based upon the 95% confidence intervals.[32] In all cases (including the case of non-normally distributed 95% confidence intervals), the larger confidence interval was used to assess statistical significance.

The statistical significance of the difference in the number of episodes produced by ‘*affiliated*’ and ‘*independen*t’ podcast series was calculated via a two-sample t-test.[33]

## 4. Results

952 science podcast series met the inclusion criteria for this study. A similar number - i.e. many hundreds of podcast series - were excluded as per the inclusion/exclusion criteria, but the details of these individual excluded podcasts were not recorded.

Between 2004 and 2010, the total number of science podcast series grew in a linear manner (see linear fit in Figure 1A, R^2^ = 0.99). In contrast, between 2010 and 2018 the total number of available science podcast series grew exponentially (see Figure 1A, R^2^ = 0.99), rising to 952 podcast series by the sampling period (5^th^ January – 5^th^ February 2018). Before 2004, 11 science podcasts were available as internet radio shows which have subsequently been made available as science podcast series.

**Figure 1:**
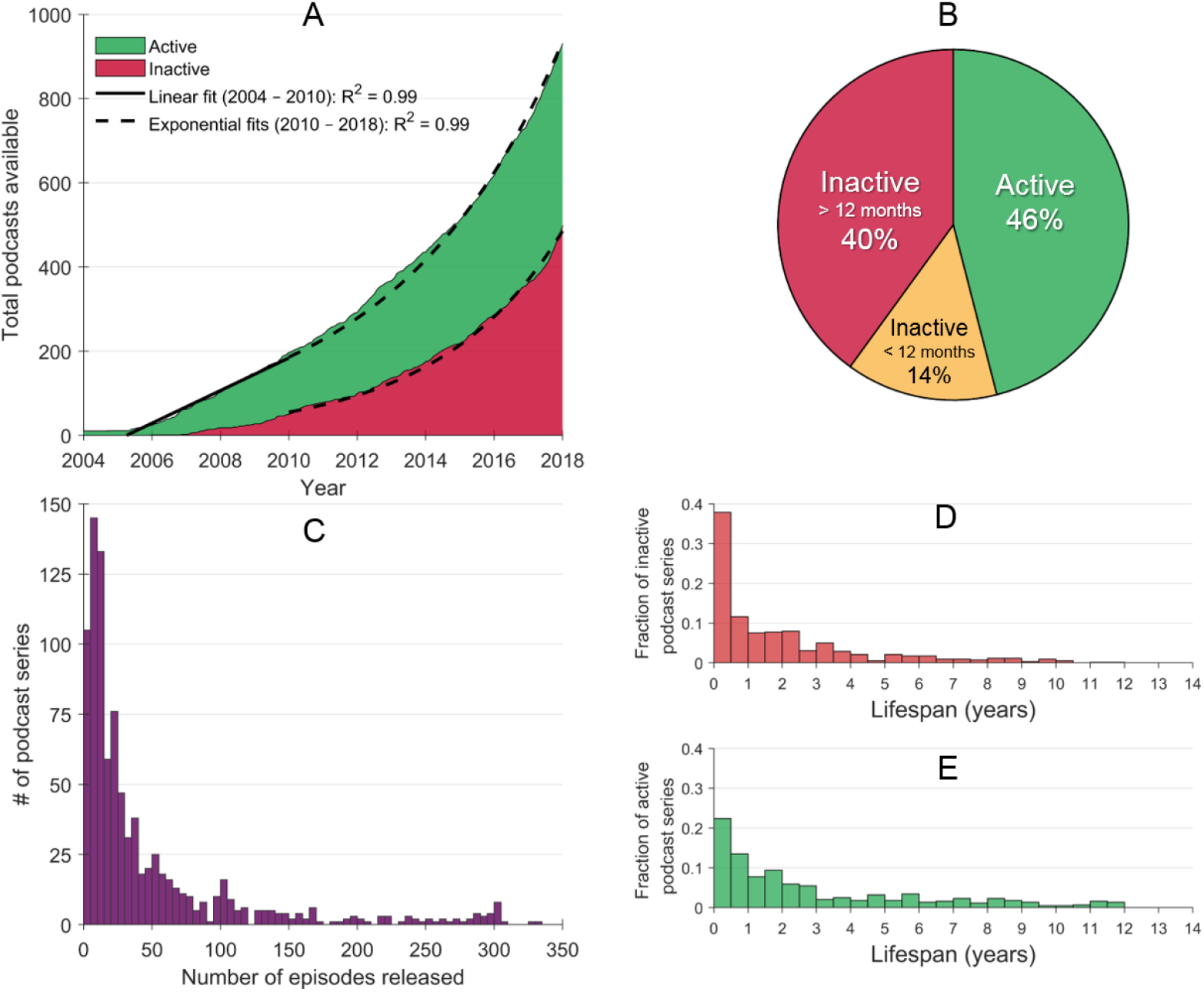
The growth and lifespan of science podcasts. **(A)** The total number of science podcasts shows linear growth between 2004 and 2010, followed by exponential growth to from 2010-2018 (n = 952). **(B)** The proportion of active/inactive science podcast series during the sampling period, i.e. between 05/01/18 and 05/02/18. **(C)** The total number of episodes released by all podcast series (NB: x-axis is constrained to 350 episodes for clarity due to outliers). **(D)** The lifespan of inactive podcasts (n = 515). **(E)** The lifespans of currently active podcasts (n = 437).

As of their individual sampling dates,^d^ 46% of total science podcast series were *‘active’*, meaning that they released an episode in the three months prior to their specific sampling date. Of the remaining ‘inactive’ podcast series, 14% released an episode between three to twelve months prior their sampling date, and 40% had not released an episode for over a year prior to their sampling date (see Figure 1B).

The number of episodes released by each science podcast series was found to be highly variable: 33% of science podcast series produced fewer than 10 episodes, and 72% of science podcast series produced fewer than 50 episodes (see Figure 1C and Table 1). From Figure 1D, it is apparent that a high proportion of science podcast series (almost 40%) did not produce podcast episodes for more than a year.

**Table 1.** The number of episodes released by science podcast series.

| Number of Episodes released | Number of Podcasts Qualifying | % |
| --- | --- | --- |
| 1 Episode | 25 | 2.6 |
| ≤ 10 Episodes | 250 | 33.0 |
| ≤ 50 Episodes | 685 | 72.0 |
| ≤ 100 Episodes | 802 | 84.2 |
| ≤ 300 Episodes | 913 | 95.9 |
| > 300 Episodes | 39 | 4.1 |
| > 500 Episodes | 17 | 1.8 |
| > 1000 Episodes | 5 | 0.5 |

| Statistical Descriptor | Number of Episodes Released<br>(entire population) |
| --- | --- |
| Modal | 10 |
| Median | 20 |
| Mean | 73 |

**Table 2.**
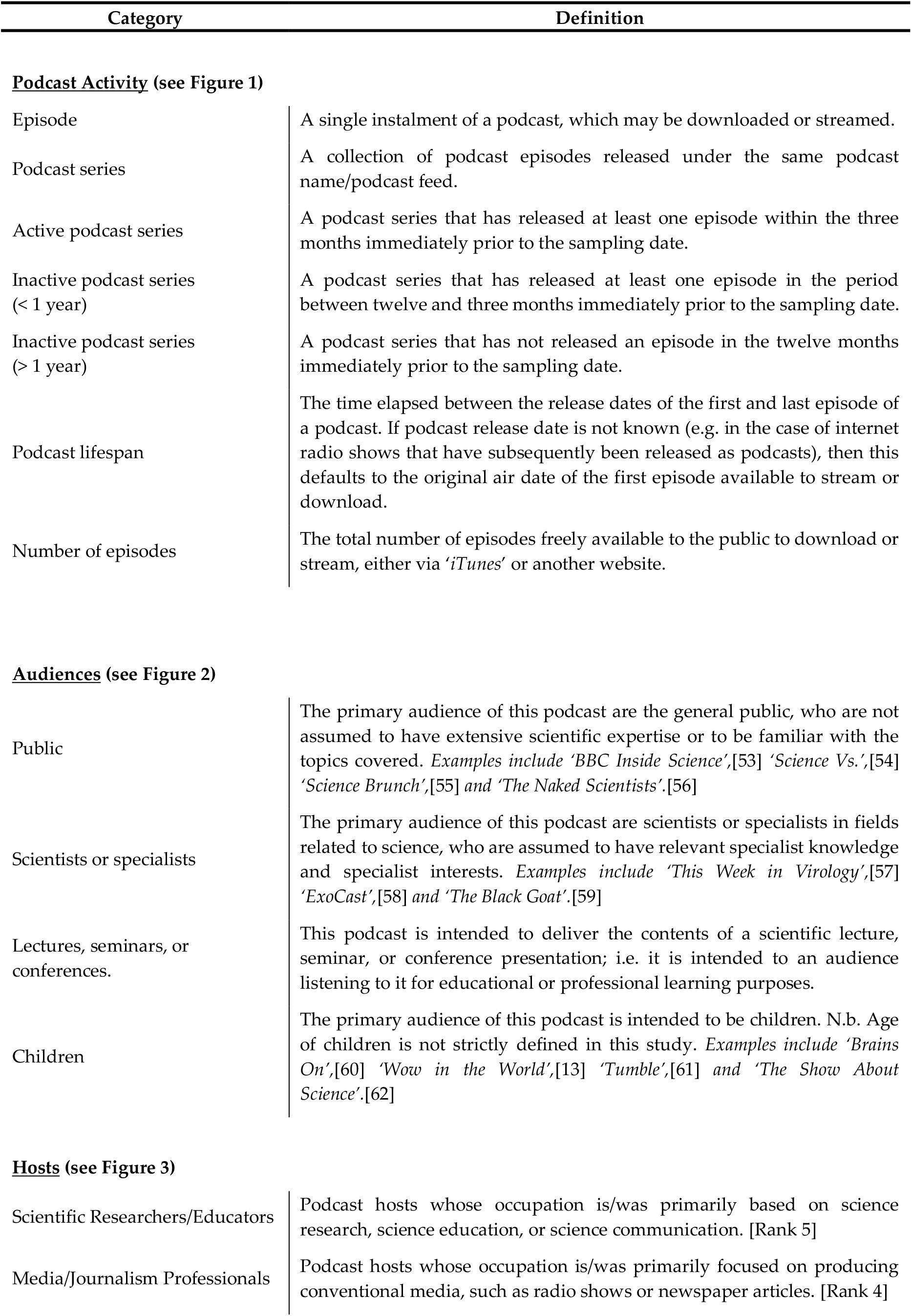
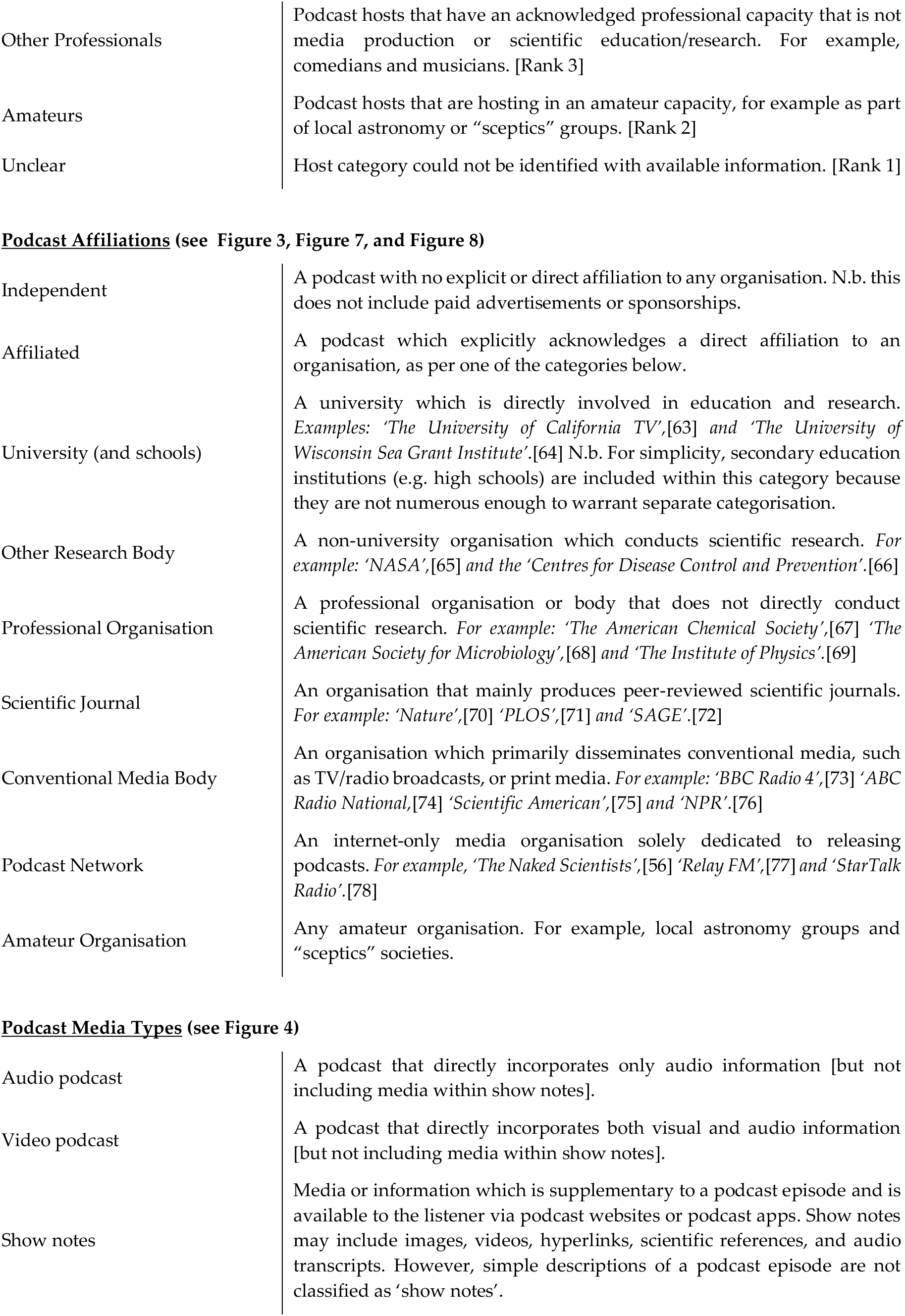
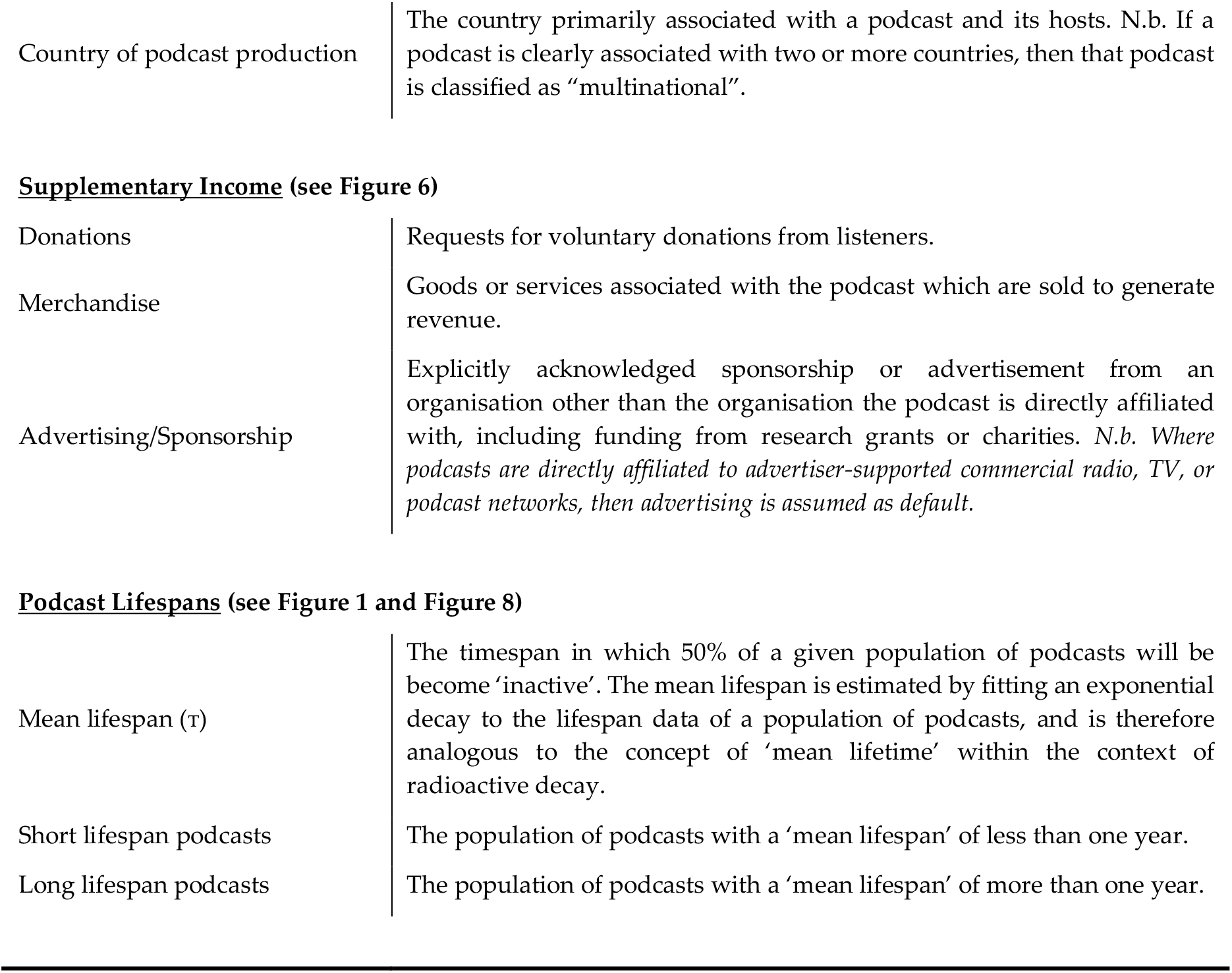
Categorical definitions used for classifying podcasts.

| Category | Definition |
| --- | --- |
| **<u>Podcast Activity</u> (see Figure 1)** |  |
| Episode | A single instalment of a podcast, which may be downloaded or streamed. |
| Podcast series | A collection of podcast episodes released under the same podcast name/podcast feed. |
| Active podcast series | A podcast series that has released at least one episode within the three months immediately prior to the sampling date. |
| Inactive podcast series (< 1 year) | A podcast series that has released at least one episode in the period between twelve and three months immediately prior to the sampling date. |
| Inactive podcast series (> 1 year) | A podcast series that has not released an episode in the twelve months immediately prior to the sampling date. |
| Podcast lifespan | The time elapsed between the release dates of the first and last episode of a podcast. If podcast release date is not known (e.g. in the case of internet radio shows that have subsequently been released as podcasts), then this defaults to the original air date of the first episode available to stream or download. |
| Number of episodes | The total number of episodes freely available to the public to download or stream, either via 'iTunes' or another website. |
| **<u>Audiences</u> (see Figure 2)** |  |
| Public | The primary audience of this podcast are the general public, who are not assumed to have extensive scientific expertise or to be familiar with the topics covered. *Examples include 'BBC Inside Science',[53] 'Science Vs.',[54] 'Science Brunch',[55] and 'The Naked Scientists'.[56]* |
| Scientists or specialists | The primary audience of this podcast are scientists or specialists in fields related to science, who are assumed to have relevant specialist knowledge and specialist interests. *Examples include 'This Week in Virology',[57] 'ExoCast',[58] and 'The Black Goat'.[59]* |
| Lectures, seminars, or conferences. | This podcast is intended to deliver the contents of a scientific lecture, seminar, or conference presentation; i.e. it is intended to an audience listening to it for educational or professional learning purposes. |
| Children | The primary audience of this podcast is intended to be children. N.b. Age of children is not strictly defined in this study. *Examples include 'Brains On',[60] 'Wow in the World',[13] 'Tumble',[61] and 'The Show About Science'.[62]* |
| **<u>Hosts</u> (see Figure 3)** |  |
| Scientific Researchers/Educators | Podcast hosts whose occupation is/was primarily based on science research, science education, or science communication. [Rank 5] |
| Media/Journalism Professionals | Podcast hosts whose occupation is/was primarily focused on producing conventional media, such as radio shows or newspaper articles. [Rank 4] |
| Other Professionals | Podcast hosts that have an acknowledged professional capacity that is not media production or scientific education/research. For example, comedians and musicians. [Rank 3] |
| Amateurs | Podcast hosts that are hosting in an amateur capacity, for example as part of local astronomy or “sceptics” groups. [Rank 2] |
| Unclear | Host category could not be identified with available information. [Rank 1] |
| **Podcast Affiliations** (see Figure 3, Figure 7, and Figure 8) |  |
| Independent | A podcast with no explicit or direct affiliation to any organisation. N.b. this does not include paid advertisements or sponsorships. |
| Affiliated | A podcast which explicitly acknowledges a direct affiliation to an organisation, as per one of the categories below. |
| University (and schools) | A university which is directly involved in education and research. *Examples: ‘The University of California TV’,[63] and ‘The University of Wisconsin Sea Grant Institute’.[64]* N.b. For simplicity, secondary education institutions (e.g. high schools) are included within this category because they are not numerous enough to warrant separate categorisation. |
| Other Research Body | A non-university organisation which conducts scientific research. *For example: ‘NASA’,[65] and the ‘Centres for Disease Control and Prevention’.[66]* |
| Professional Organisation | A professional organisation or body that does not directly conduct scientific research. *For example: ‘The American Chemical Society’,[67] ‘The American Society for Microbiology’,[68] and ‘The Institute of Physics’.[69]* |
| Scientific Journal | An organisation that mainly produces peer-reviewed scientific journals. *For example: ‘Nature’,[70] ‘PLOS’,[71] and ‘SAGE’.[72]* |
| Conventional Media Body | An organisation which primarily disseminates conventional media, such as TV/radio broadcasts, or print media. *For example: ‘BBC Radio 4’,[73] ‘ABC Radio National’,[74] ‘Scientific American’,[75] and ‘NPR’.[76]* |
| Podcast Network | An internet-only media organisation solely dedicated to releasing podcasts. *For example, ‘The Naked Scientists’,[56] ‘Relay FM’,[77] and ‘StarTalk Radio’.[78]* |
| Amateur Organisation | Any amateur organisation. For example, local astronomy groups and “sceptics” societies. |
| **Podcast Media Types** (see Figure 4) |  |
| Audio podcast | A podcast that directly incorporates only audio information [but not including media within show notes]. |
| Video podcast | A podcast that directly incorporates both visual and audio information [but not including media within show notes]. |
| Show notes | Media or information which is supplementary to a podcast episode and is available to the listener via podcast websites or podcast apps. Show notes may include images, videos, hyperlinks, scientific references, and audio transcripts. However, simple descriptions of a podcast episode are not classified as ‘show notes’. |
| **Countries** (see Figure 5) |  |
| Country of podcast production | The country primarily associated with a podcast and its hosts. N.b. If a podcast is clearly associated with two or more countries, then that podcast is classified as “multinational”. |
| **Supplementary Income** (see Figure 6) |  |
| Donations | Requests for voluntary donations from listeners. |
| Merchandise | Goods or services associated with the podcast which are sold to generate revenue. |
| Advertising/Sponsorship | Explicitly acknowledged sponsorship or advertisement from an organisation other than the organisation the podcast is directly affiliated with, including funding from research grants or charities. *N.b. Where podcasts are directly affiliated to advertiser-supported commercial radio, TV, or podcast networks, then advertising is assumed as default.* |
| **Podcast Lifespans** (see Figure 1 and Figure 8) |  |
| Mean lifespan ($\tau$) | The timespan in which 50% of a given population of podcasts will be become ‘inactive’. The mean lifespan is estimated by fitting an exponential decay to the lifespan data of a population of podcasts, and is therefore analogous to the concept of ‘mean lifetime’ within the context of radioactive decay. |
| Short lifespan podcasts | The population of podcasts with a ‘mean lifespan’ of less than one year. |
| Long lifespan podcasts | The population of podcasts with a ‘mean lifespan’ of more than one year. |

A wide variety of science podcast series topics/themes were recorded, with 66% of science podcast series themed around discipline-specific topics (see Figure 2A). Of particular note, *‘Chemistry’* was the topic for only 3% of science podcast series, compared to 18% for *‘Physics and Astronomy’*, and 14% for *‘Biology’*. 34% of science podcast series were categorised as *‘General Science’*, i.e. science podcasts focusing on no single discipline-specific theme.

**Figure 2:**
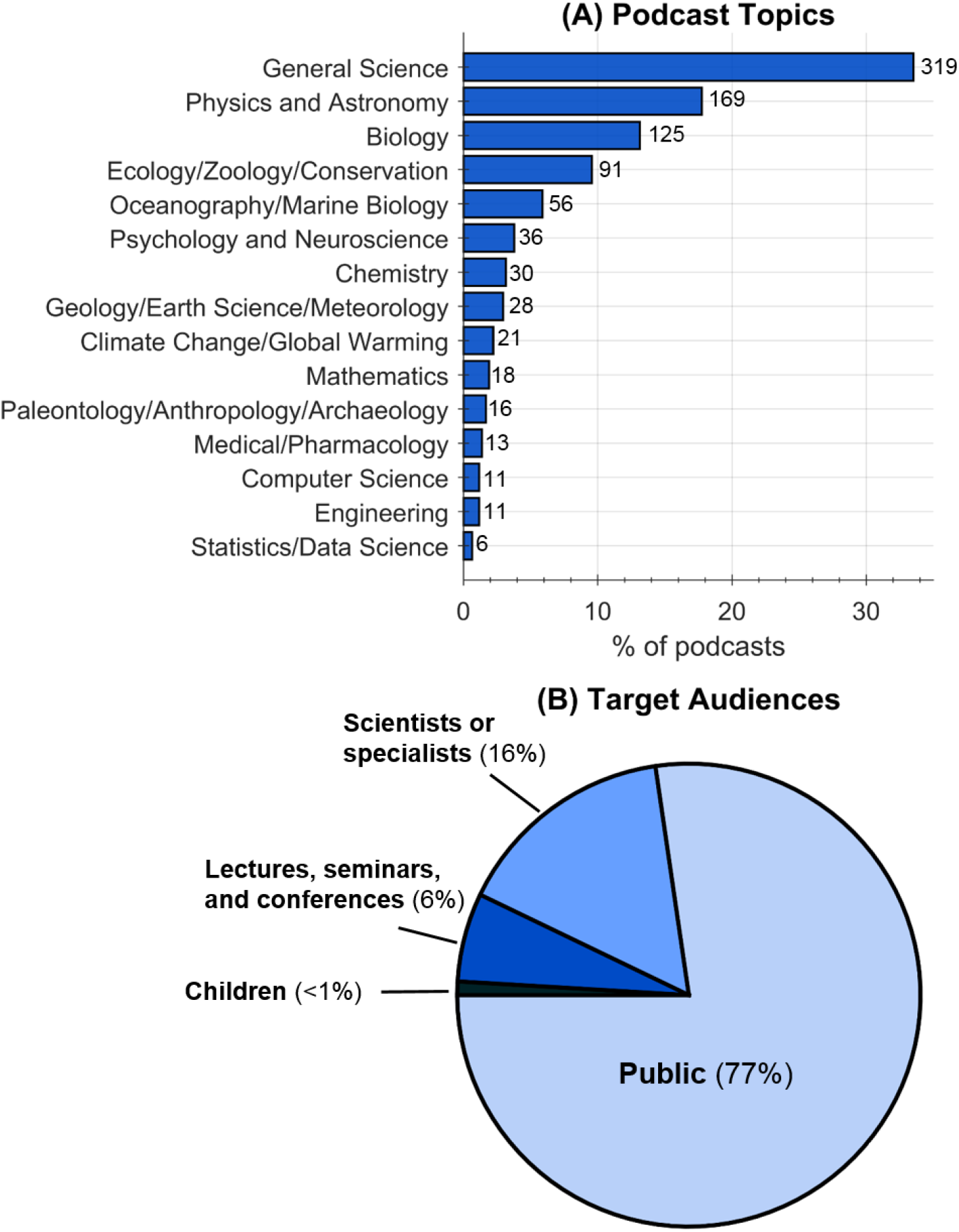
What are the scope and aims of science podcasts? **(A)** The proportion of science podcasts dedicated to various scientific topics. **(B)** The target audiences of science podcasts.

The majority of science podcast series (77%) have been targeted to public audiences, 16% were targeted towards scientists or specialists, and 6% were provided as academic lectures, research seminars/conferences, or as secondary education learning aids (see Figure 2B).

Nearly 2/3^rds^ (65%) of science podcast series were hosted by ‘*scientists*’; 10% were hosted by *‘media professionals’*, 7% by *‘other professionals’*, and 5% by *‘amateurs’* (see Figure 3A). Host categories could not be identified for 13% of science podcast series.

**Figure 3:**
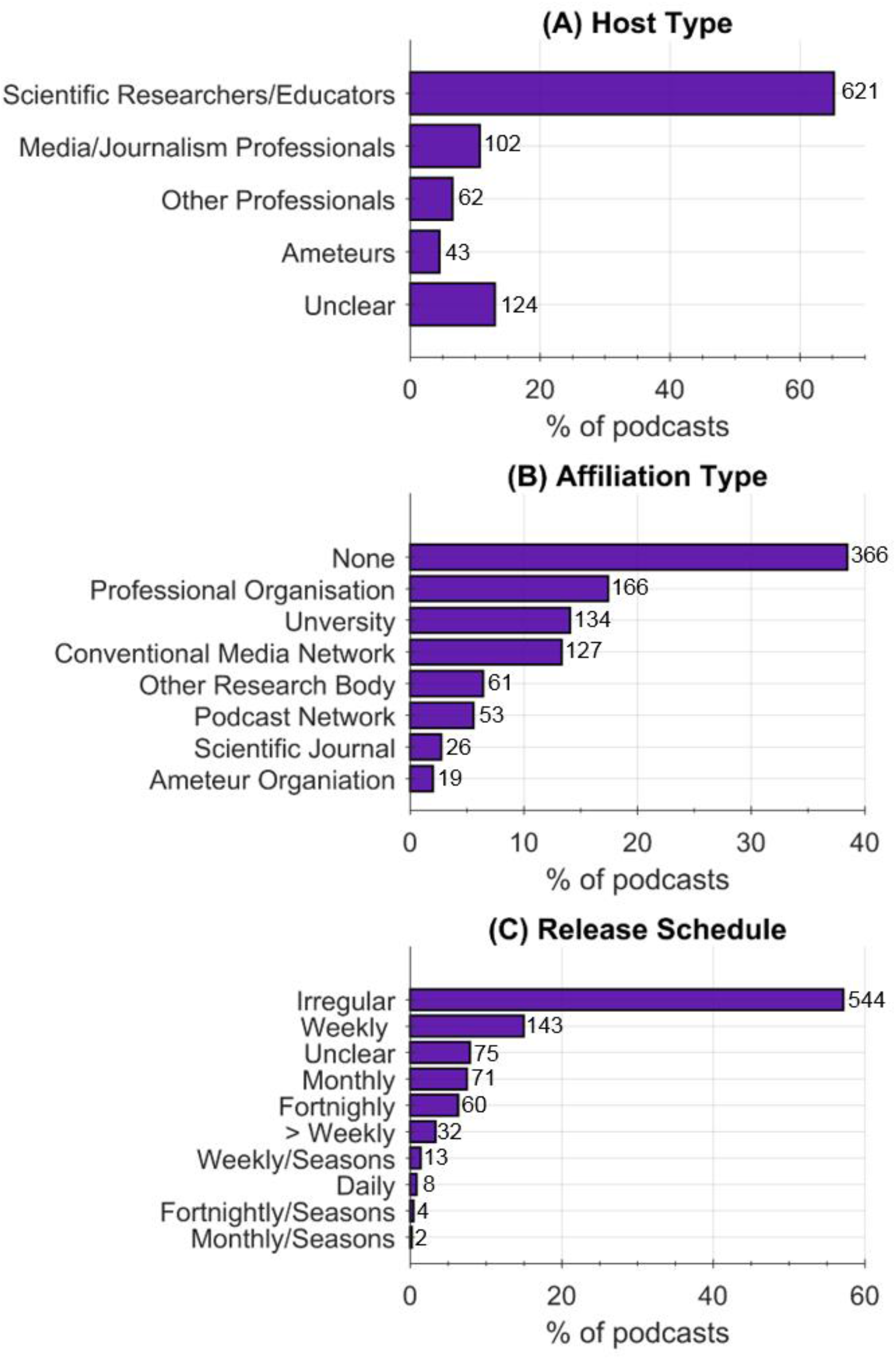
Who produces science podcasts? **(A)** The backgrounds of science podcast hosts. **(B)** The organisational affiliations of science podcasts. **(C)** The release schedule of science podcasts.

38% of science podcast series were produced independently, and 62% were produced with some explicitly acknowledged affiliation to an organisation (see Figure 3B). *‘Professional Organisations’* produced 17% of science podcasts; *Universities’* 14%; *‘Conventional Media Networks’* 13%; *‘Other Research Bodies’* 6%; *‘Podcast Networks’* 5%; *‘Scientific Journals’* 3%, and *‘Amateur Organisations’* 2%. How podcast affiliation, or lack thereof, affects various science podcast production outputs is explored further, later in this manuscript.^e^

57% of science podcast series did not follow a regular episode release schedule (see Figure 3C). The most popular release schedule was *‘Weekly’* (15%), followed by *‘Monthly’* (8%), and *‘Fortnightly’* (6%). Only 3% of science podcasts released more than one episode per week, and 1% released an episode daily. Only 2% of science podcast series explicitly acknowledged a seasonal release format, i.e. periods of scheduled episode releases followed by an extended period where no episodes are released.

Whilst podcasts can contain both audio and visual information, 87% of science podcast series were audio-only, with the remaining 13% being video podcast series (so-called *“vodcasts”*) (see Figure 4A). 51% of science podcast series provided additional non-audio supplementary material in the form of show notes (e.g. hyperlinks, images, references, etc.) (see Figure 4B). From Figure 4C, it is clear that the proportion of new video science podcast series produced each year, as a fraction of overall science podcast series, has declined from a peak of ~30% of science podcast series in 2007 to ~5% of science podcast series in 2017. However, the absolute number of new video science podcast series produced each year has been relatively constant, at around 9 ± 3 (mean ± standard deviation). This long-term decline in video podcasts may reflect changing behaviour, i.e. that audiences consume podcasts whilst undertaking activities incompatible with watching video content.[3–5,22]

**Figure 4:**
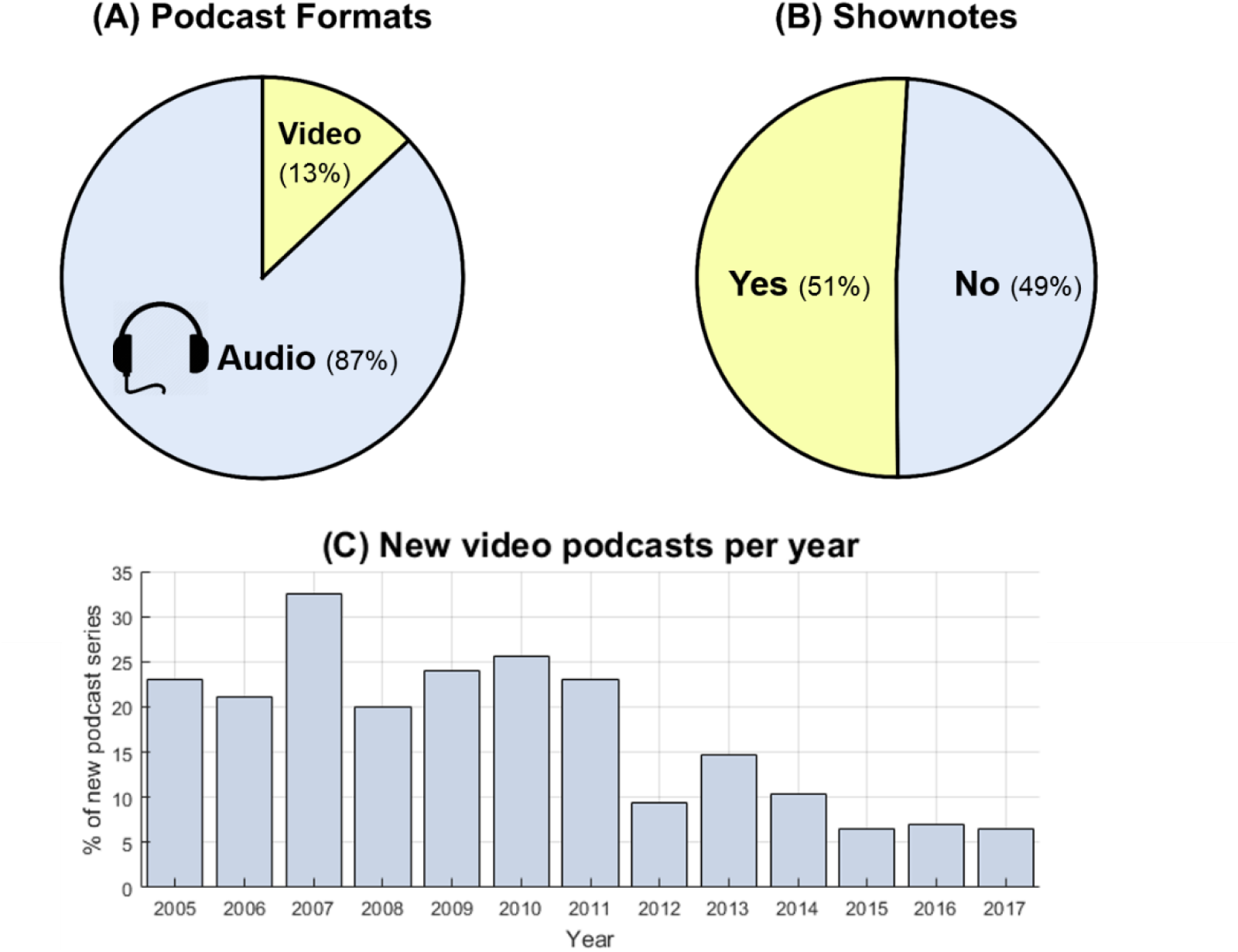
non-audio media in science podcasts. **(A)** The proportion of audio-only science podcasts compared to video format science podcasts. **(B)** The usage of show notes by science podcasts. **(C)** New video science podcasts produced each year as a proportion of the overall number of science podcasts produced each year. Long term declines in the number of video podcasts produced can be seen.

Global production of science podcast series to date is shown in Figure 5: 57% of the available English language science podcast series were produced in the United States of America (USA); 17% were produced in the United Kingdom (UK); 5% in Australia; 3% in Canada, and 1% in the Republic of Ireland. Other countries produce a combined total of 7% of English language science podcast series. A country of production could not be identified for 10% of science podcast series.

**Figure 5:**
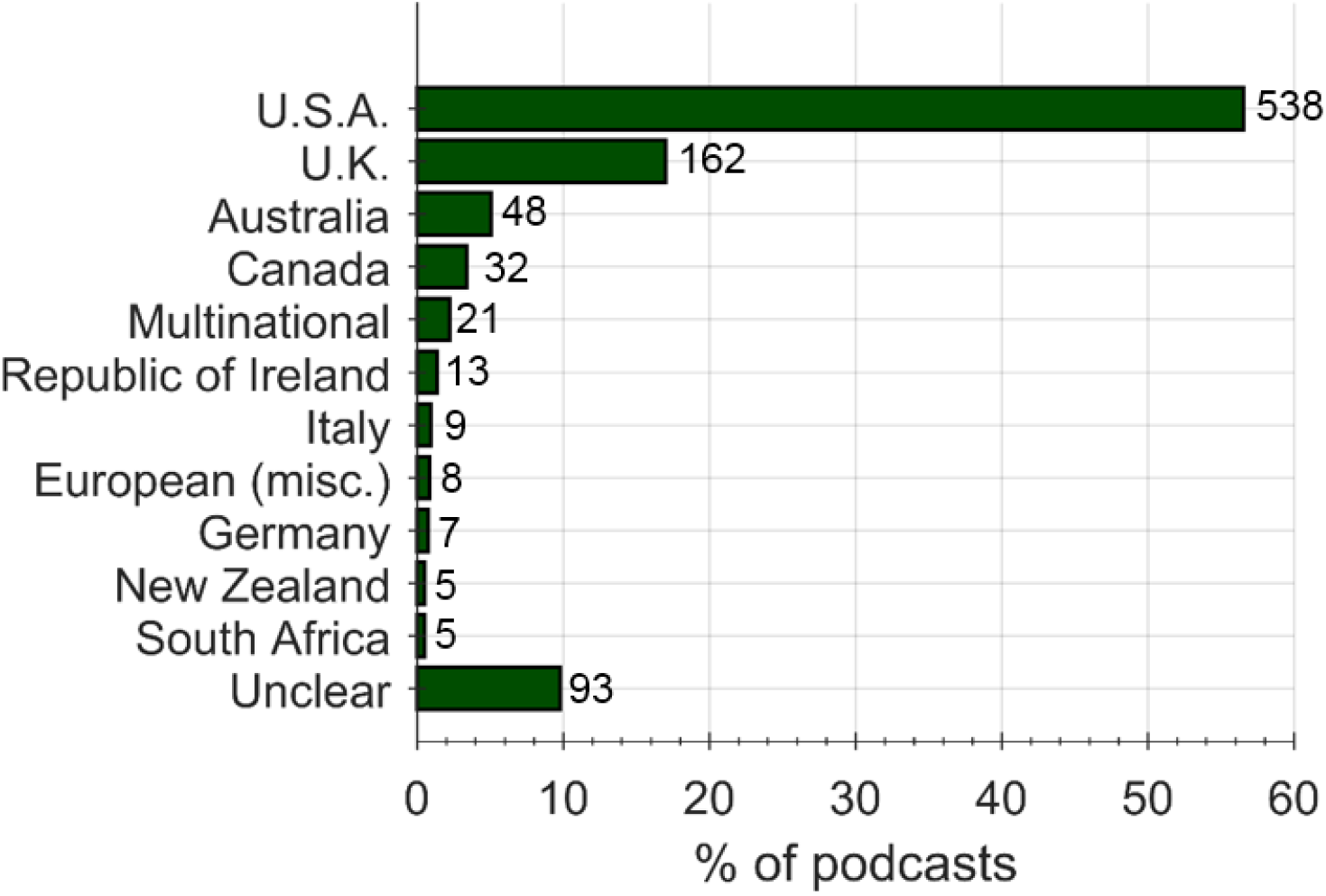
Production of English language science podcasts by country.

76% of science podcast series were observed to have no overt supplementary income mechanisms and are thus seemingly independently financed by their producers (see Figure 6A). *‘Advertising’* was the least commonly utilised supplementary income mechanism (see Figure 6B), but it was common for science podcasts to mix *‘Voluntary Donations’, ‘Merchandise’*, and *‘Advertising’* to various degrees.

**Figure 6:**
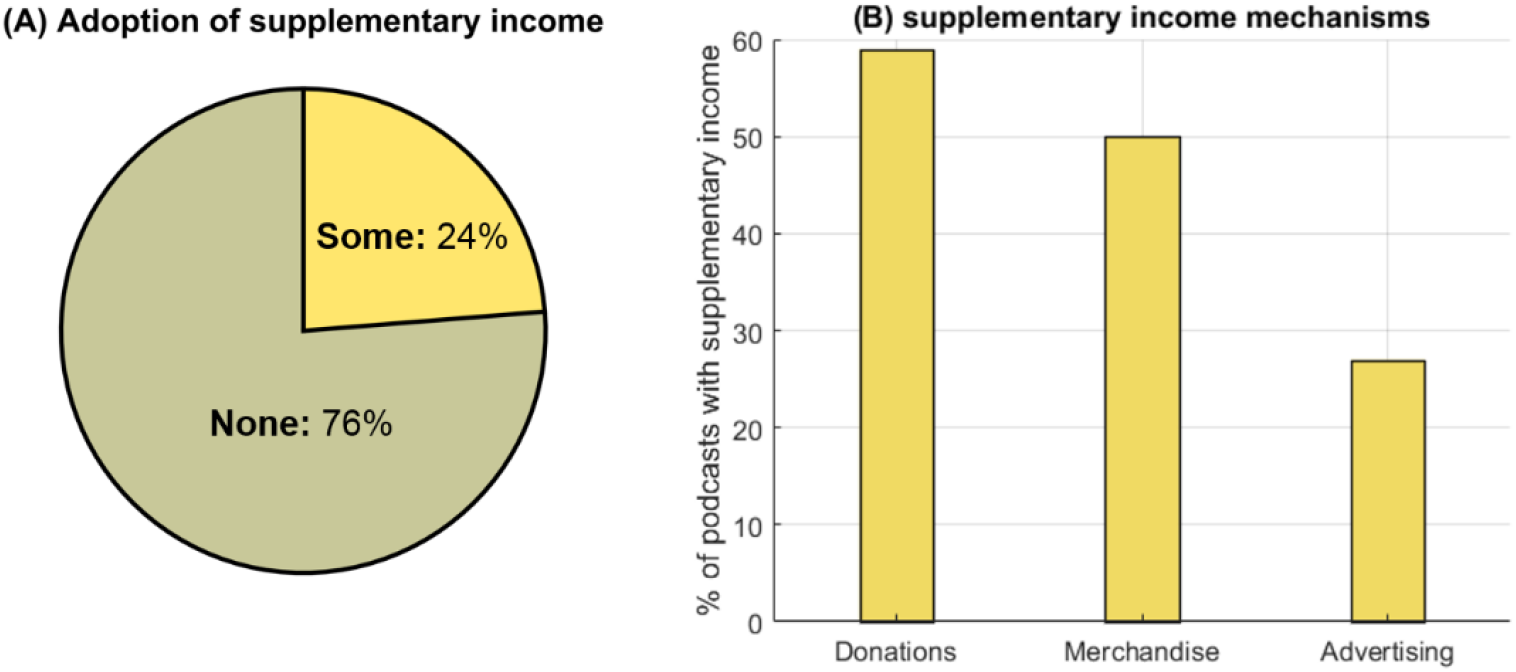
Do science podcasts generate overt supplementary income? **(A)** The proportion of podcasts with some supplementary income mechanism vs the proportion that have none. **(B)** The percentage of the subset of science podcasts with a supplementary income, that use each type of supplementary income mechanism. N.b. these categories are not mutually exclusive as some science podcasts utilise multiple income mechanisms.

The differences between ‘*independent*’ science podcast series and ‘*affiliated*’ science podcast series in relation to various production outputs is shown in Figure 7. In terms of podcast activity, there is only a marginal difference between the percentage of active ‘*affiliated*’ and ‘*independent*’ science podcast series (48% and 45% respectively) (see Figure 7A). However, a larger proportion of *‘independent’* podcast series (84%) are targeted to the public, compared to *‘affiliated’* podcast series (73%) (see Figure 7B). A slightly smaller proportion of *‘independent’* podcast series (14%) are targeted towards *‘scientist/specialist’* audiences compared with *‘affiliated’* podcast series (17%) (see Figure 7B). Nearly all science podcast series billed as academic seminars, student lectures, or secondary education aids are produced as *‘affiliated’* podcast series (see Figure 7B). Roughly 75% of both *‘independent* and *‘affiliated’* podcast series had no overt supplementary income (see Figure 7C). However, a considerably greater proportion of *‘independent’* podcast series solicited for *‘voluntary donations’* and sold *‘merchandise’* (see Figure 7C). *‘Advertising’* was much more prevalent for *‘affiliated’* podcast series (25%) than *‘independent’* podcast series (11%) (see Figure 7C); this is likely due to many ‘*affiliated*’ podcast series being associated with commercial broadcast networks, where ‘*advertising*’ was assumed.

**Figure 7:**
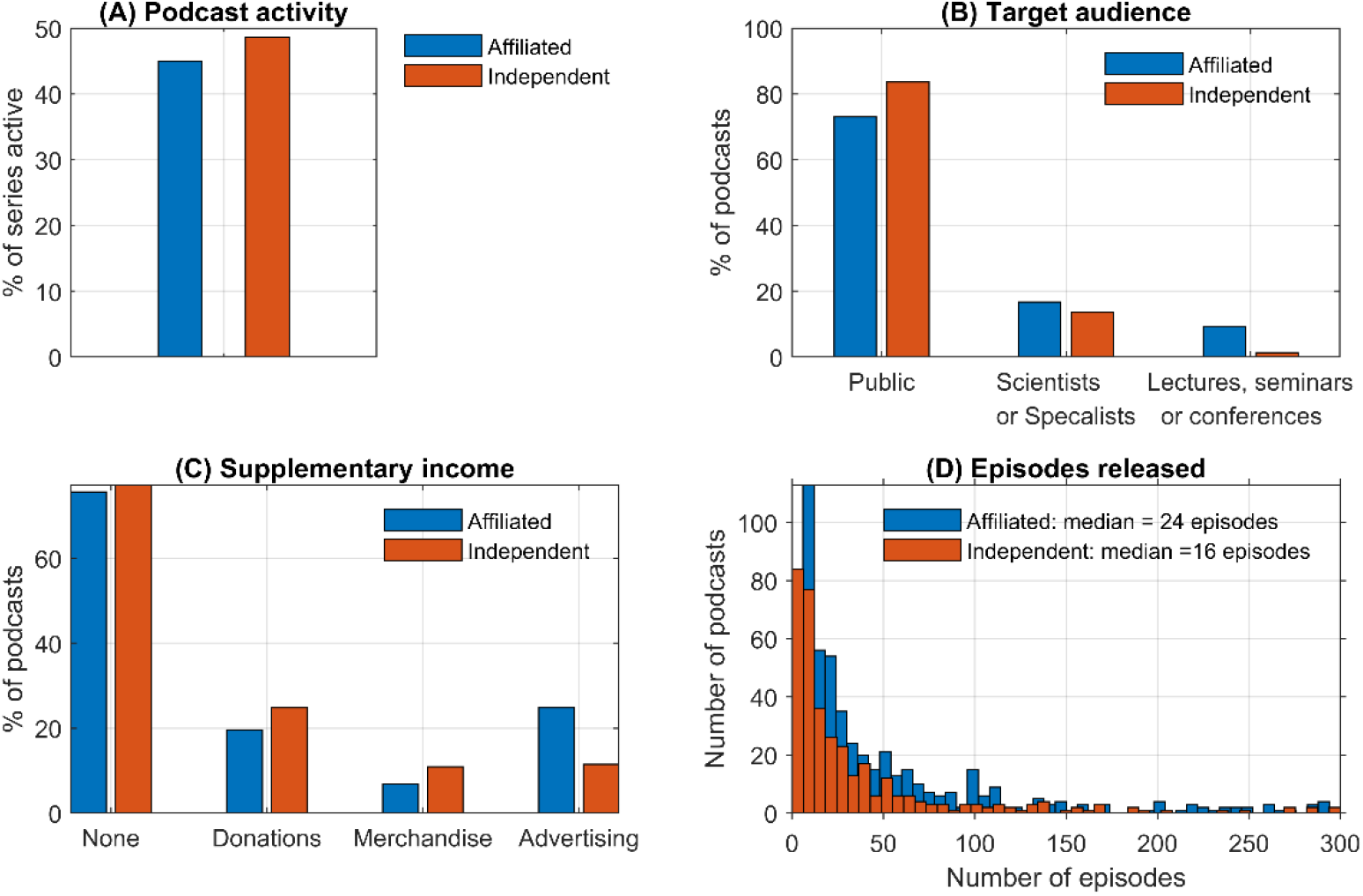
Does science podcast affiliation alter production outcomes? **(A)** Podcast affiliation vs. % of active podcasts. **(B)** Podcast affiliation vs. target audience. **(C)** Podcast affiliation vs. supplementary income mechanisms. **(D)** Podcast affiliation vs. total number of podcast episodes produced by podcast series, showing that affiliated podcasts produce a greater number of episodes (median = 24, average = 48) than independent podcasts (median 16, average = 90) (p < 0.01).

‘*Affiliated*’ podcast series produced a greater number of podcast episodes (median = 24, average = 90), than *‘independent’* podcast series (median = 16, average = 48). A two-tailed t-test found that the difference between in the overall number of episodes released was statistically significant (p = 0.01) and that the greater average number of podcast episodes released by *‘affiliated’* podcast series was also statistically significant (p < 0.01)

The lifespan of both *‘independent’* and *‘affiliated’* podcast groupings was best-fitted by a two-term exponential. This indicates that both ‘affiliated’ and ‘independent’ podcast groupings contain subsets of *‘short lifespan’* and *‘long lifespan’* podcast series (see Figure 8A and Figure 8B). Extraction of fit parameters enables the estimation the podcast *‘mean lifespan’* (T) for each of these podcast subsets. T is analogous to the concept of *‘mean lifespan’* in radioactive decay; i.e. T is the elapsed time span in which, 50% of the podcasts in a population become inactive. The best-fit and 95% confidence interval values for T are shown in Figure 8C and Figure 8D. For short-duration podcast series subsets, the difference in the best-estimates of T for *‘affiliated’* and *‘independent’* podcast series was not statically significant (p>0.33). However, for long-duration podcast series subsets, the difference in the best-estimates of T or *‘affiliated’* and *‘independent’* podcast series (5.5 years, and 4.3 years respectively) was statistically significant (p < 0.02).

**Figure 8.**
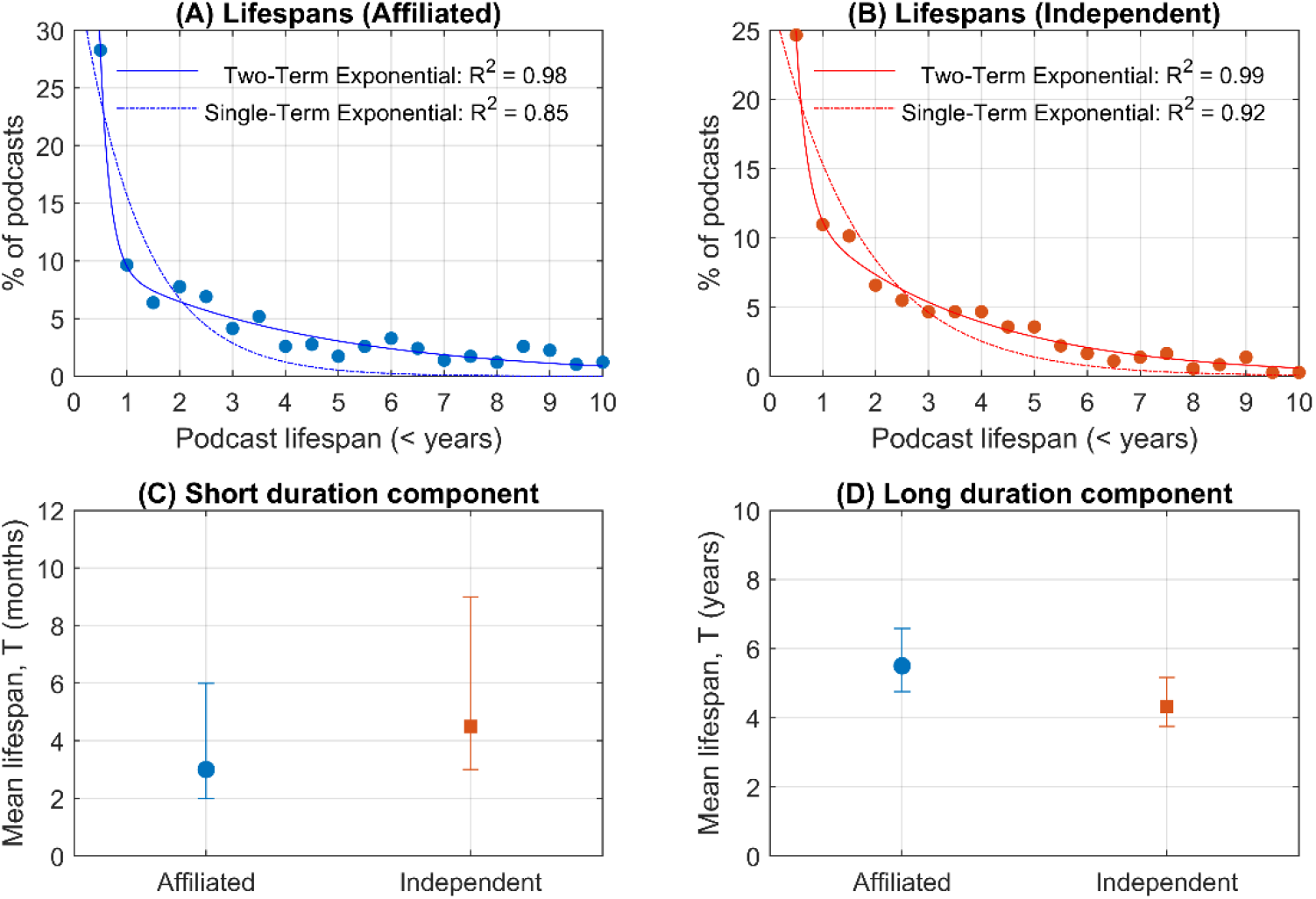
Estimated mean lifespans of podcasts. **(A)** two-term exponential fit to the lifespan of *‘affiliated’* podcasts. **(B)** Two-term exponential fit to the lifespan of ‘*independent*’ podcasts. **(C)** Mean lifespans of short-duration podcast estimated from the two-term exponential fits. Points represent the best-fit estimate and error bars represent 95% confidence intervals. The difference between best-estimate values is not statistically significant. **(D)** Mean lifespans of long-duration podcasts estimated. Points represent the best-fit estimate and error bars represent 95% confidence intervals. The difference between best-estimate values was statistically significant [p < 0.02].

## 5. Discussion

### Methodology and associated limitations

This is the first study to analyse the global production and outputs of a large group of science podcast series. As such, the findings here provide fundamental and novel insight into who is producing science podcast series and their target audiences. However, before detailed discussion of results, it is important to acknowledge the limitations of the methodology employed in this study.

Firstly, in this study, only English language science podcast series were surveyed and analysed. It is highly probable that non-English language science podcast series would demonstrate different trends due to different listener and producer demographics.

Secondly, it is important to note that the data generated in this study was analysed (coded) by only a single researcher (the author). This is a shortcoming of the study design because different individuals may categorize qualitative data different. Best practice in such research would have been to follow “multiple coding” procedures, i.e. for multiple researchers to evaluate and analysing the data, subsequently resolving any discrepancies arising, whilst also maximising robustness in data coding.[34] Also relevant to data coding and interpretation off the results is that a host classification based on a notional ranking of scientific authority was used. The rationale of this system was that having even a single scientist in a podcast host group will tend to elevate the scientific content of a podcast, therefore such instances should be highlighted. However, this host classification system has several limitations: (1) it is based on analysis of textual and visual data, (2) it may overly-simplify the data in a manner that over-represents higher-ranked host classifications (i.e. scientists and media professionals), and (3) it doesn’t consider the expertise of guests on podcasts. For future studies, a classification system that better represents the myriad possibilities of podcast host backgrounds should be implemented.

Thirdly, science podcast series were primarily identified by survey of only a single *‘iTunes’* category: i.e. the *‘Natural Sciences’* category.[31] This is similar to the methodology of a previous study by Birch and Weitkamp, which defined science podcasts as *“the natural sciences and mathematics”*.[15] However, constraining this study to the *‘Natural Sciences’* category limits the podcasts examined for two reasons: (1) listing a podcast on *‘iTunes’* is not mandatory; (2) the category a podcast as listed on *‘iTunes’* is self-selected by the uploader, and therefore, many science podcasts may have been listed in *‘iTunes’* categories not examined. The most obvious category that wasn’t analysed was the *‘Science and Medicine’* category.[35] However, a large number of podcast series that covered dubious/harmful pseudo-medical practices and advice were prevalent within the *‘Science and Medicine’* category. Therefore, an extremely stringent and in-depth inclusion/exclusion criteria strategy would have to be developed and applied, along with deep content analysis (e.g. actually listening to individual episodes of each podcast), to ensure that only legitimate scientific podcast series are included in any such study. Unfortunately, this was beyond the scope of the current study. Moreover, some science podcast series are not listed on *‘iTunes’* at all; an example of such a science podcast is *‘BioLogic Podcast’*, which is hosted on the video sharing website *‘YouTube’*.[36] Additionally, it should be noted that some podcast series may voluntarily restrict the number of podcast episodes that are freely available to the public via ‘*iTunes*’ or other websites, but only freely-available episodes were included for analysis within this study. Therefore, this study provides a *lower-bound* on the number of science podcast series available during the sampling period.

Fourthly, this study exclusively examined the visual and textual online presence of podcast series. Due to practical constraints, it was not possible to examine the extensive audio data associated with science podcasts. Therefore, it is possible that various aspects of podcast production were not fully categorised. This could affect all studied podcast categories, but most likely affects the capture of any audio-only advertisements or sponsorships that were not acknowledged in textual or visual web content of science podcasts. Therefore, it is possible that a greater proportion of science podcasts contain advertisements or sponsorships than is explicitly reported by this study. With regards to hosts, it is possible that podcasts hosts and production teams fit multiple categories, but this is not capture by the relatively shallow nature of our study; as Picardo and Regina (2008)[8] note in their detailed comment on podcasting: “defining who is inside and who is outside [sic: the podcast] control room is not an easy task”.

Fifthly, podcast episode length data and podcast download statistics were not available for analysis. Such data would be desirable for a more complete analysis of analysis of the consumption and production of science podcasts.

A notable limitation of this study is that the original podcast upload date for radio shows broadcast pre-2004 are not known; instead the original air-date episodes (as provided on iTunes or another relevant website) is used as a compromise. This accounts for the 11 podcast series available prior to 2004 (see supplementary database for full details). Of these 11 podcast series, 10 are affiliated to an organisation. Considering that 586 *‘affiliated’* podcast series were analysed and that the mean lifespan, T, is calculated from robust curve-fitting models, the influence of these 10 podcast series on the results of lifespan fitting calculations can be considered negligible for the purposes of this study.

### Science podcasts vs. general podcasts

Large-scale studies of podcast production have not been published in peer-reviewed literature, therefore it is necessary to look beyond the peer-reviewed literature to glean large-scale podcast production insights. In 2015, Morgan published a semi-formal study of podcasts of many different topics as a blog post on ‘ *medium.com’*.[27] Whilst not published in a peer-reviewed journal, all data associated with Morgan’s study is publicly available. Morgan’s study sampled a subset of podcast series available on *‘iTunes’* in June 2015. Morgan estimated that there were 206,000 unique podcast series available on *‘iTunes’* at that time. Morgan than selected a random subset of podcast series for further analysis. This subset consisted of a total of 2500 podcast series, with 100 random podcast series drawn from the 25 “most popular” *‘iTunes’* categories (N.B. this did not include any category theme around science). Morgan’s sampling and analysis was fully-automated, so manual categorisation of podcast production outputs was not conducted. Importantly, Morgan defined *“active podcast series”* as podcast series that had released an episode within the 6 months prior to the sampling date [27]; this is a less stringent definition than that used in the present study, which defines *“active podcast series”* as podcast series that had released an episode within 3 months prior to the sampling date. Morgan found that the number of podcast series available on *‘iTunes’* had grown from ~10,000 in 2007 to ~206,000 in 2015. When graphed, the trends in growth of total number of podcast series calculated by Morgan (not shown here) appear broadly similar to the trends shown in Figure 1A, i.e. displaying distinct linear growth up to 2010, and exponential growth thereafter. This indicates that trends in the growth of science podcast series likely reflects the overall growth of the podcast medium. Additionally, Morgan found that roughly 40% of podcast series were *‘active’* by his less stringent definition.[27] This is lower than the comparable population of *‘active’* science podcast series (46%) found by the present study (see Figure 1B). This comparison suggests that science podcast series may be more inclined to continue to release episodes compared to the wider population of podcast series. However, this comparison may not necessarily be valid because Morgan did not exclude podcast series that had not released a single episode. Further, Morgan found that the average lifespan of podcast series was around 6 months, and that podcasts, on average, released 12 episodes, at a rate of 2 episodes per month. Additionally, Morgan estimated that around 20% of podcast series listed on ‘*iTunes*’ at the time were not English language podcasts.

### Insights into the production of science podcasts

The predominance of scientists as hosts for science podcast series (see Figure 3A), combined with fact that most science podcast series (57%) are released on an irregular schedule (see Figure 3C), may indicate that a significant majority of science podcast series are being produced by scientists as an extra commitment beyond their regular duties as a scientific researcher, science educator, or science communicator. However, the limitations of the study methodology must be considered in that this study may possibly over-represent scientists as podcast hosts (see the Discussion sub-section ‘Methodology and Associated Limitations’). The result that most science podcasts do not have any overt supplementary income mechanisms (see Figure 5A) is of note when considering that there can be substantial costs associated with hosting a podcast (i.e. high-quality audio equipment and editing software, as well as branded websites for advertisement and podcast hosting). The lack of overt supplementary income mechanisms suggests that independent science podcast hosts are paying these costs “out of their own pocket”. These results combine to give a broad impression that many science podcast series are being produced by scientists with no financial recompense. The obvious exception being the science podcast series *‘affiliated’* to organisations that can provide undisclosed financial support. However, the fundamental validity of this interpretation requires further research and study before firm conclusions can be made.

Figure 2A shows that only 3% of science podcast series cover ‘*chemistry*’ as their main topic. When compared to the two other primary science subjects typically taught in schools - i.e. *‘biology’* (13% of science podcast series), and *‘physics and astronomy’* (18% of science podcasts) – it appears that chemistry is under-represented in science podcasts. There are several potential explanations as to why this may be. A 2011 editorial in the journal *‘Nature Chemistry’* suggested that chemistry *“is a central science”*, meaning that aspects of chemistry are incorporated into other disciplines (e.g. biochemistry and materials research); therefore chemistry is often not distinctly represented in public-facing science communication.[37] Similarly, Hartings and Fahly (2011) noted that popular science involving chemistry may not be labelled as chemistry; that chemistry is complex; and that chemistry lacks unifying themes and public narratives that may be present in biology and physics.[38] Additionally, a review of chemistry communication in 2016 noted that concepts in chemistry are well-served by dynamic visual representations,[39] therefore chemistry may not be well-suited to the primarily-audio format of podcasts. Indeed, chemistry content is very well received in more visual internet mediums, e.g. the video series: *‘Periodic Videos’* on *‘YouTube’*.[40] Velden and Lagoze (2009) note that chemistry has been slow to adopt “new web-based models of scholarly communication” when compared to physics and biology.[41] Whilst this may true for scholarly communications, it is not clear if this is true for chemistry and digital science communication practices. All these reasons are likely to play into the apparent lack of chemistry science podcast series. This reinforces a 2016 recommendation from the *‘National Academies of Science, Engineering, and Medicine’*, that science funding agencies should support digital media for chemistry communication as a priority.[42]

The statistically significant greater best-estimate values for mean lifespan of ‘*affiliated*’ podcast series (5.5 years) compared to *‘independent’* podcast series (4.3 years) (see Figure 8D) could be explained by the hypothesis is that ‘i*ndependent*’ podcast series may be more likely to be produced by individuals or small groups, with limited time and resources, whereas ‘*affiliated*’ podcast series are produced by organisations with dedicated staff with defined duties. Such dedicated staff could take-over podcasting duties when necessary, therefore extending the overall lifespan of the ‘*affiliated*’ podcast series compared to ‘*independent*’ podcast series. However, no firm conclusions with regards to the causes of podcast series sustainability can be drawn from this study, and it should be noted that there are exceptionally long-running podcast series within both the ‘*independent*’ and ‘*affiliated*’ subsets. In their 2011 study titled *“Why podcasters keep going”*, Markman found that creator-audience community, engagement (e.g. via emails, discussion forums, social media etc), audience appreciation, and enjoyment were key drivers of podcast longevity. Markman notes that further study is required into the phenomena of podcast longevity and so-called *“podfading”*, where podcasts are no longer produced.[43]

### Open questions and future directions

This study provides the first large-scale overview of the production of English language science podcast series, yet there are many open questions that remain. For example, does the general content of science podcasts differ across different cultures and languages?[10] What level of prior knowledge is required to understand science podcasts?[44] Are science podcasts helping to change non-representative stereotypes of scientists?[45] Do science podcasts promote and foster trust in science?[16] Are podcasts considered in long-term science communication and impact strategies?[46]

The motivations for podcast hosts and creators for podcast have previously been explored in two studies: Markmann (2011)[43], and Markman and Sawyer (2014).[17] However, the motivations for the creation of science podcast series may be rather different from the motivations of podcast producers for other topics. For example, how do factors such as career recognition (or lack thereof), and time constraints motivate science podcasters,[47] and how do podcast creators use social media to engage with their audiences?[48]

In recent years, new methods of analysis have been developed for other new online media such as blogs and online news sources.[44,49] Whilst metrics such as listener numbers and attention are not available for large-scale analysis of podcasts, other techniques could be adapted to the study of science podcasts. For example, analysis of hyperlinks included in blogs has been used to provide a measure of “content diversity”.[49] Similarly, hyperlink analysis could be applied to science podcast show notes to ascertain diversity of sources and content that audiences are referred to.

Audiobooks are an increasingly popular medium [50] that could be used as a direct comparison between the written word and audio forms of science communication. Audiobooks, like podcasts, are a portable and convenient audio-only format. Audiobooks are typically narrated by a single voice-actor or by the author themselves. However, because they are typically direct adaptions of the written word, science audiobooks are formal, not conversational.[51] A further distinction of audiobooks from podcasts is that audiobooks are nearly exclusively produced by for-profit media and publishing companies, not independent, decentralised, content creators. As an example of the potential richness of audiobooks as a data source: at the time of writing, ‘*Audible*’, (a major for-profit audiobook content provider), has over 2000 science audiobooks available across ‘*science*’, ‘*astronomy*’, ‘*physics’, and biology’* categories.[52] Therefore, audiobooks could serve as a “test-bed” for studies comparing how media formats may alter the effectiveness of science communication.

## 6. Conclusions

This study has revealed large-scale trends in science podcasting for the first time. Overall, the total number of science podcast series grew linearly between 2004 and 2010, and subsequently it has grown exponentially between 2010 and 2018. A total of 952 science podcast series met the inclusion criteria for this study, giving a lower-bound on English language science podcasts available at the start of 2018. Most science podcast series (87%) are audio-only, with the number of new video-format science podcast series declining from a peak of ~30% in 2007 to only 5% in 2017. This may reflect that podcast audiences are choosing to listen to podcasts whilst undertaking activities incompatible with consuming video content.

One third of science podcast series were found to cover many aspects of science, but many individual subjects were well represented by dedicated podcast series. Notably, ‘chemistry’ as a topic appears to be under-represented, with only 3% of podcast series compared to 18% for ‘physics and astronomy’, and 13% for ‘biology. This apparent under-representation in podcasting may mirror similar long-term trends in science communication where chemistry has been under-represented as a distinct subject. This may also reflect the idea that chemistry is best-represented by visual mediums, i.e. not audio podcasts.

Most science podcasts appear to be targeted towards the audience of the general public (77%), with fewer science podcast series serving educational purposes (6%), serving specialist audiences (16%), or dedicated to science communication for children (< 1%). 51% of science podcast series included extra information to audiences in the form of supplementary show notes, containing text, images, or hyperlinks.

Almost 2/3rds of science podcast series have at least one host with a background in scientific research, science communication, or science education. This indicates that scientists are using podcasts to communicate with the public. The exact reasons as to why podcasting is attractive to science communicators are still to be ascertained, but it is likely to be due to the simplicity of producing podcasts, the low amount of equipment required, the global audience reach, the ability to receive feedback via social media, the intimate nature of the medium, and the lack of format constraints.

38% of science podcast series appeared to be produced independently; the remaining 62% of science podcast series had an overt affiliation to some sort of organisation, e.g. a university, funding agency, or media network. Generally, most science podcast series appeared to not have any overt form of supplementary income, i.e. through advertising, selling merchandise, or soliciting for audience donations. This indicates that a large portion of science podcast series are being financed by independent content creators or by organisations. Of podcasts with overt supplementary income, podcasts *‘affiliated* with an organisation were more likely to have adverts, and *‘independent’* science podcast series were more likely to sell merchandise or solicit for audience donations. Whether or not a science podcast series is independent or affiliated to an organisation appears to make key differences in several production outputs. Most notably, *‘independent’* podcast series produce fewer episodes on average (median 16, average 48) than *‘affiliated’* podcast series (median 24, average 90) [p ≤ 0.01]. Furthermore, the long-term mean-lifespan of *‘independent’* podcasts (4.3 years) appears to be significantly less than the long-term mean-lifespan of *‘affiliated’* podcasts (5.5 years) [p < 0.02].

Whilst this study has provided the first insights into the large-scale production of science podcasts, there are still many ongoing questions about how science podcasts are being used to communicate science. Metrics for download and listener attention were not available for the podcasts studied, but content analysis of show-note hyperlinks could be used in future as a proxy for content diversity. Audiobooks could serve as a medium for comparative studies between written and spoken science communication, without the conversational nature of podcasts. In future, a combination of quantitative and qualitative approaches may be required to yield further insights into the motivations of science podcasters, why they choose to produce the podcasts that they do, and how science podcasts are meeting the need for science communication without geographic barriers.

## Supporting information

## Data Accessibility

Supporting data available on BioRxiv at: https://doi.org/10.1101/298356

## Acknowledgements

Thanks to the many science podcast creators who inspired this study. Thanks to C.A. Osnes for proof-reading of draft manuscripts.

## Footnotes

a Note that the term ‘podcast’ can both refer to a single podcast episode or a series of podcast.

b *‘iTunes’* may also be referred to elsewhere as *‘Apple Podcasts’*.[79]

c Two-term exponential fits were necessary because single-term exponential decays were found to fit the data poorly, as shown in Figure 8.

d The exact sampling date for each podcast is provided in the associated supplementary dataset.

e See Figure 7 and Figure 8

